# *Rickettsia* Sca2 Recruits Two Actin Subunits for Nucleation but Lacks WH2 Domains

**DOI:** 10.1101/467431

**Authors:** SS Alqassim, IG Lee, R Dominguez

## Abstract

The *Rickettsia* ~1,800 amino acid autotransporter protein Sca2 promotes actin polymerization on the surface of the bacterium to drive its movement using an actin comet tail mechanism. Sca2 mimics eukaryotic formins in that it promotes both actin filament nucleation and elongation and competes with capping protein to generate filaments that are long and unbranched. However, despite these functional similarities, Sca2 is structurally unrelated to eukaryotic formins and achieves these functions through an entirely different mechanism. Thus, while formins are dimeric, Sca2 functions as a monomer. However, Sca2 displays intramolecular interactions and functional cooperativity between its N- and C-terminal domains that are crucial for actin nucleation and elongation. Here, we map the interaction of N- and C-terminal fragments of Sca2 and their contributions to actin binding and nucleation. We find that both the N- and C-terminal regions of Sca2 interact with actin monomers, but only weakly, whereas the full-length protein binds two actin monomers with high affinity. Moreover, deletions at both ends of the N- and C-terminal regions disrupt their ability to interact with each other, suggesting that they form a contiguous ring-like structure that wraps around two actin subunits, analogous to the formin homology-2 (FH2) domain. The discovery of Sca2 as an actin nucleator followed the identification of what appeared to be a repeat of three WH2 domains in the middle of the molecule, consistent with the presence of WH2 domains in most actin nucleators. However, we show here that contrary to previous assumptions Sca2 does not contain WH2 domains, and that the corresponding region is folded as a globular domain that cooperates with other parts of the Sca2 molecule for actin binding and nucleation.

## Introduction

Numerous pathogens, including several bacteria and some viruses, take advantage of the actin cytoskeleton of the host eukaryotic cell for motility and infection (1-3). A common mechanism relies on the activation of actin nucleation on the surface of pathogens to generate actin comet tails that propel their movement (4). Pathogens that use this mechanism typically nucleate actin by exposing proteins that mimic eukaryotic nucleation promoting factors (NPFs), which recruit and activate the host Arp2/3 complex (5). However, some bacteria encode their own actin nucleators, i.e. proteins capable of directly activating the polymerization of eukaryotic actin, including *Vibrio* VopL/F (6-9), *Burkholderia* BimA (10) and *Rickettsia* Sca2 (11-13). Such bacterial nucleators have evolved independently of their eukaryotic counterparts, and they tend to polymerize actin using different molecular mechanisms than the eukaryotic nucleators (4). This feature, along with the requirement of actin-based motility for infection, highlights the importance of studying bacterial nucleators for a better understanding of both pathogenicity and actin nucleation mechanisms.

Here, we focus on Sca2 (surface cell antigen 2), a surface protein of spotted fever *Rickettsiae*. *Rickettsiae* are intracellular gram-negative pathogenic bacteria that produce severe human diseases such as typhus and spotted fever (14). Upon infection of eukaryotic cells, they undergo actin comet tail-based motility by using two different proteins expressed on the surface of the bacterium at different stages of infection. Early post infection, RickA, a protein that mimics eukaryotic NPFs, recruits the host Arp2/3 complex to activate the formation of actin comet tails that are short and curved (15-17). Later post infection, Sca2, a protein that mimics eukaryotic formins, polymerizes actin directly, creating actin comet tails that are long and unbranched (11, 12, 17)

Sca2 is an autotransporter protein. Like other autotransporters, Sca2 uses an N-terminal signal sequence (residues 1-33 in *R. conorii* Sca2, the species studied here) to cross the inner bacterial membrane, after which it inserts a C-terminal translocator domain (residues 1516-1795) into the outer membrane to form a pore through which the rest of the protein (passenger domain, residues 34-1515) is translocated and exposed on the surface of the bacterium (18, 19). The requirement to pass through the narrow translocation pore and fold on the outside of the bacterium in the absence of an energy source imposes substantial constraints on the folding of autotransporter proteins. As a consequence, most autotransporters display relatively simple folds, based on a repeating basic folding unit, which in most cases consists of β-strands that stack up to form a β-solenoid (20, 21).

The structure of Sca2 residues 34-400, which we previously determined (12), suggests that Sca2 constitutes an exception among autotransporters in that the basic folding unit consists of helix-loop-helix motifs, which stack upon one another to give rise to an all-helical structure. We refer to this fragment of Sca2 as the N-terminal repeat domain (NRD), because a structural superimposition of the various helix-loop-helix motifs revealed conserved amino acids at specific locations. There is no high-resolution structural information for the remaining ~1,115 amino acids that form the passenger domain of Sca2. However, helix-loop-helix repeats of the kind observed in the structure of NRD are clearly discernable at the sequence level between residues 1181 and 1341, which we thus refer to as the C-terminal repeat domain (CRD). Moreover, the entire passenger domain is predicted to be primarily helical, as experimentally confirmed for a fragment comprising residues 868-1515 using circular dichroism (12). Based on these considerations, we have proposed that most of passenger domain of Sca2 adopts the same basic fold observed in the structure of NRD (12). This poses a problem, because toward the middle of the passenger domain (residues 868-1023) Sca2 presents what is thought to be a repeat of three Wiskott-Aldrich syndrome homology 2 (WH2) domains. Consistent with this idea, simultaneously mutating all three of the predicted WH2 domains within full-length Sca2 impairs the nucleation activity (12). However, repeats of WH2 domains do not form part of globular structures in any of the proteins analyzed thus far; instead WH2 domains are typically connected by flexible linkers of ~15 amino acids (22). In Sca2 the separation between the predicted WH2 domains is much greater, 45-55 amino acids, and the intervening regions are folded. Here we show that, contrary to previous assumptions, Sca2 does not have WH2 domains, and that the 868-1023 region cooperates with other parts of the Sca2 protein for actin binding and nucleation.

Sca2 is also unique among bacterial nucleators in that it functions by a formin-like mechanism (11, 12). Like eukaryotic formins, Sca2 promotes both actin nucleation and elongation, i.e. it remains bound to filament barbed ends after nucleation and drives the processive incorporation of profilin-actin while also competing with capping protein (11, 12). Like formins, Sca2 generates long, unbranched filaments. Yet, despite these functional similarities, Sca2 is structurally unrelated to formins, and performs these functions through an entirely different mechanism (12). Also unlike formins, which form dimers, Sca2 is monomeric. However, it displays intramolecular interactions and cooperativity between its N- and C-terminal halves that are crucial for actin nucleation and elongation (12). Here, we map the interaction of N- and C-terminal fragments of Sca2 and their contributions to actin binding and nucleation and conclude that the two fragments interact in a head-to-tail circular manner to recruit two actin subunits for nucleation, somewhat analogous to the formin homology-2 (FH2) domain.

## Materials and Methods

### Protein expression and purification

All the constructs used in this study were amplified from the *Rickettsia conorii* sca2 gene (GenBank: AAL02648). All of the constructs, except Sca_1090-1355_, were expressed and purified as described previously (12), with slight modifications. Briefly, constructs Sca_34-1515_, Sca_34-670_, Sca_34-400_, Sca_421-670_, Sca_869-1060_, Sca_1090-1515_, and Sca_868-1515_ were cloned into the pTYB11 vector (New England BioLabs) containing a chitin affinity tag and intein domain. Sca_1090-1355_ was cloned into the pRSFduet vector with an N-terminal His-tag and an engineered N-terminal TEV protease site. All the proteins were expressed in *E. coli* Rosetta or Arctic Express cells grown in Terrific Broth media. Cells were resuspended in Chitin buffer (20 mM HEPES pH 7.5, 500 mM NaCl, 1 mM EDTA, 4 mM Benzamidine) or, in the case of construct Sca_1090-1355_, nickel lysis buffer (20 mM Tris-HCl pH 7.5, 200 mM NaCl, 20 mM Imidazole, 4 mM Benzamidine) and lysed using a microfluidizer (MicroFluidics Corporation). After purification on a chitin affinity column, the chitin tag was removed by self-cleavage of the intein, induced by the addition of 50 mM DTT for 1-2 days at 4°C or 20°C, which leaves no extra residues on the Sca2 constructs. For Sca_1090-1355_, the N-terminal His-tag was cleaved after elution from the Nickel affinity column by the addition of TEV protease at 4°C overnight, followed by a second round of Nickel affinity to remove the TEV protease and any uncut protein. All of the proteins were additionally purified by either ion exchange chromatography or size exclusion chromatography. Point mutations in the WH2 domains of Sca_34-1515_ were introduced using the QuikChange II XL site-directed mutagenesis kit (Stratgene) as before (12), and expressed and purified as described above.

The predicted WH2 peptides of Sca2 (Sca_871-890_, Sca_942-962_, Sca_1003-1022_) and that of WAVE1 were cloned as MBP (Maltose-Binding Protein) fusions into a modified pMAL-c2x vector containing an N-terminal His-tag and an engineered N-terminal TEV protease site, by primer-extension to express proteins with the following overall organization: His-TEV-MBP-linker(AAANNNNNNNNNNLG)-WH2. These fusion proteins were expressed in *E. coli* Arctic cells, and purified by Nickel affinity followed by ion exchange chromatography.

### Analytical gel filtration chromatography

To assess for the formation of complexes between the various N- and C-terminal domains of Sca2, proteins were mixed, incubated for 30 minutes, and run through a Superose 6 10/300 GL column (GE Healthcare) in a buffer containing 20 mM Tris-HCl pH 7.5, 200 mM NaCl, 1 mM DTT. Peak fractions were analyzed by SDS-PAGE.

### Actin polymerization assay

Actin polymerization was measured as the fluorescence increase resulting from the incorporation of pyrene-labeled actin into filaments, using a Cary Eclipse Fluorescence Spectrophotometer (Varian). Before data acquisition, 2 μM Mg-ATP-actin (6% pyrene-labeled) was mixed with different concentrations of Sca2 constructs in F-buffer (10 mM Tris-HCl pH 7.5, 1 mM MgCl_2_, 50 mM KCl, 1 mM EGTA, 0.1 mM NaN_3_, 0.2 mM ATP). Data acquisition started 5-10 s after mixing. All of the measurements were done at 25°C. Control experiments were carried out with the addition of buffer alone.

### Isothermal titration calorimetry

ITC measurements were carried out on a VP-ITC instrument (MicroCal). Protein samples were dialyzed for 2 d against the ITC buffer (20 mM HEPES pH 7.5, 200 mM NaCl, 1 mM DTT, and 35 μM LatB when monomeric actin is present). Titrations consisted of 10 μl injections, lasting for 25 s, with an interval of 300 s between injections. The heat of binding was corrected for the heat of injection, determined by injecting proteins into buffer. Data were analyzed using the program Origin (OriginLab). The temperature and parameters of the fit (stoichiometry and affinity) of each experiment are given in the figures.

## Results

### Intramolecular interaction of the N- and C-terminal regions of Sca2

Previously, we had shown that the N- and C-terminal regions of Sca2, fragments 34-670 (Sca_34-670_, see Fig. 1A for construct definition) and 670-1515 (Sca_670-1515_), interact with each other, i.e. they co-purified when co-expressed, and together they replicate the activity of the full-length passenger domain (Sca_34___1515_) (12). Furthermore, deleting ~200 amino acids near the middle of the protein (residues 671-867) did not significantly affect this interaction nor the nucleation activity; co-expressed constructs His-Sca_34-670_ and GST-Sca_868-1515_ elute together during purification and their complex can reproduce most of the nucleation activity of Sca_34-1515_ (12). To better understand this intramolecular interaction and the overall organization of Sca2 domains for nucleation, we tested the effect of N- and C-terminal truncations of constructs Sca_34-670_ and Sca_863-1515_ on the interaction. The two interacting constructs Sca_34-670_ and Sca_868-1515_ were used as a control in this analysis. Because, previously we had tested their interaction only through co-purification using two different affinity tags (12), here we expressed the Sca_34-670_ and Sca_868-1515_ fragments independently and free of tags (see Material and Methods), and characterized their interaction quantitatively using isothermal titration calorimetry (ITC). Fitting of the ITC titration of Sca_34-670_ (in the syringe) into Sca_868-1515_ (in the ITC cell) showed that the two fragments interact with relatively high affinity (K_D_ = 7.0 ± 0.3 μM) and the reaction was exothermic in character (Fig. 1B). Furthermore, when mixed together, the two independently purified Sca2 fragments run as a single peak containing approximately equal amounts of both proteins by analytical gel filtration and SDS-PAGE analysis (Fig. 1C), consistent with the formation of a relatively tight complex.

**Figure 1.**
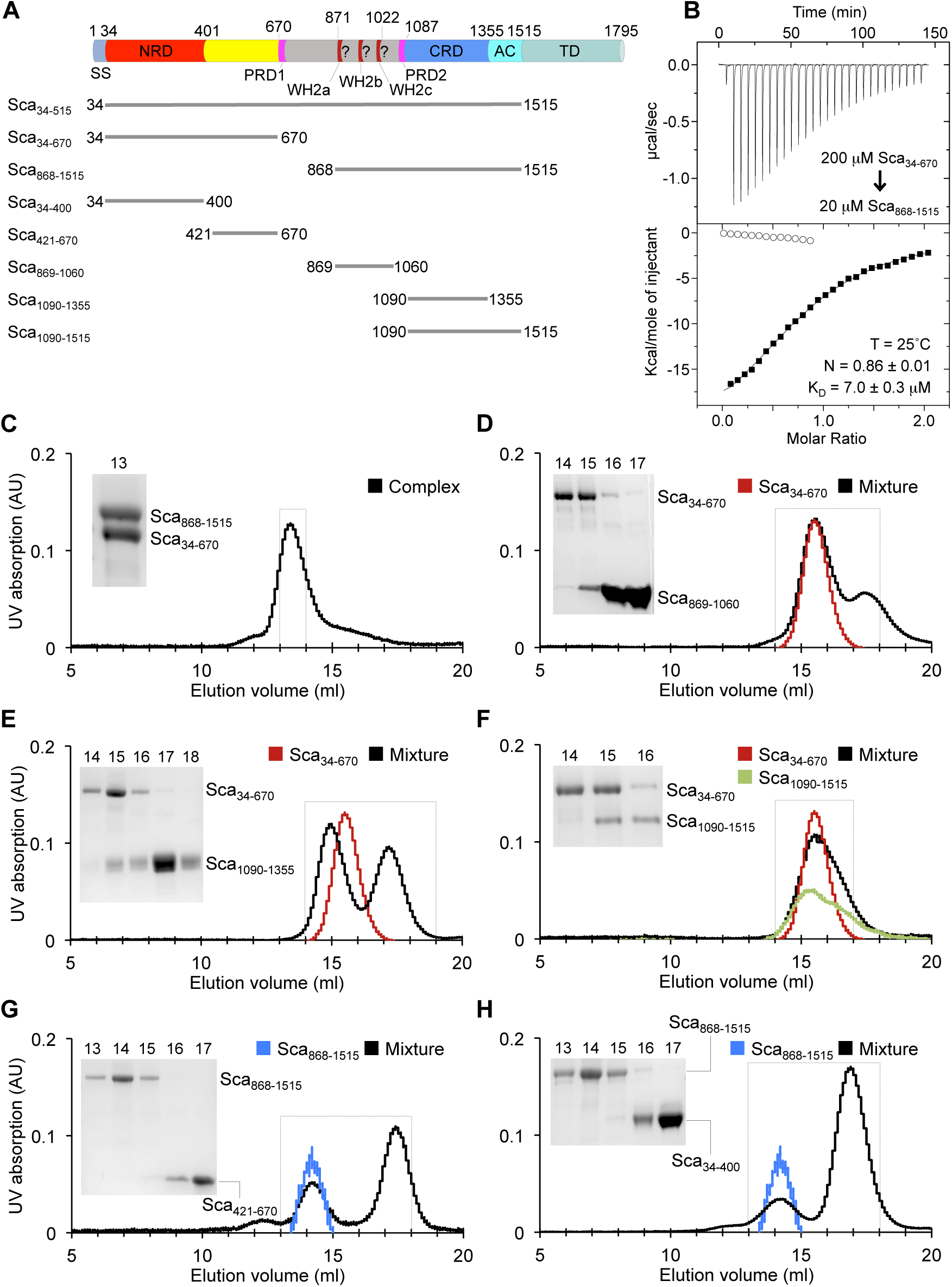
Intramolecular interaction of the N- and C-terminal regions of Sca2. (A) Domain organization of Sca2 and constructs used in this study (SS, signaling sequence; NRD, N-terminal repeat domain; PRD1,2, proline-rich domains; WH2a-c, putative WH2 domains; CRD, C-terminal repeat domain; AC, predicted autochaperone domain; TD, translocator domain).
(B) ITC titration of 200 μM Sca_34-670_ (in the syringe) into 20 μM Sca_868-1515_ (in the cell). The experiment was performed at 25°C. The dissociation constant (K_D_) and binding stoichiometry (N) derived from fitting of the binding isotherm are listed. Errors correspond to the s.d. of the fits. Open symbols correspond to titrations into buffer.
(C-H) Analytical size exclusion chromatography (SEC) and SDS-PAGE analyses of mixtures of untagged N- and C-terminal Sca2 constructs as indicated. The gels shown as insets correspond to the SEC fractions of the peak indicated by a dashed box. Part C shows that the premixed constructs Sca_34-670_ and Sca_868-1515_ run together by SEC, consistent with the formation of a complex. Parts D-F show that the N-terminal fragment Sca_34-670_ (red trace) runs separately from N- and C-terminal deletions of construct Sca_868-1515_ when premixed (black trace), consistent with lack of interaction. Parts G and H show that the C-terminal fragment Sca_868-1515_ (blue trace) runs separately from N- and C-terminal deletions of constructs Sca_34-670_ when premixed (black trace), consistent with lack of interaction.

To identify the minimal regions necessary for this intramolecular interaction, we tested binding of the intact N-terminal fragment (Sca_34-670_) with independently-expressed N- and C-terminal truncations of the C-terminal fragment (Sca_868-1515_), and *vice versa* (Fig. 1 D-H). Our results show that truncations at either end of the N-terminal (Sca_34-400_ and Sca_421-670_) or C-terminal (Sca_869-1060_, Sca_1090-1355_ and Sca_1090-1515_) regions of Sca2 abrogate the interaction, since the pre-mixed proteins emerge separately by analytical gel filtration and SDS-PAGE analysis of the peak fractions. For one construct combination, Sca_34-670_ and Sca_1090-1355_, the gel filtration peak of the individually purified proteins partially overlaps with that of their mixture, but SDS-PAGE analysis shows the two constructs are largely segregated at either end of the gel filtration peak, consistent with lack of interaction (Fig. 1F). We conclude that the minimal region necessary for the intramolecular interaction of Sca2 consists of fragments Sca_34-670_ and Sca_868-1515_. Moreover, these two fragments seem to interact in a circular, head-to-tail manner, since deletions at either end of both constructs interferes with their ability to form a complex.

### Sca2 binds two actin subunits with high affinity

The FH2 domain of eukaryotic formins also interacts in a circular, head-to-tail manner to form a dimeric ring-like structure, which encircles in the middle two actin subunits at the barbed end of the actin filament (23, 24). Each subunit of the FH2 dimer contributes to the overall binding affinity, which is significantly higher for the dimer (K_D_ ~ 2.9 μM) (25-27). We hypothesized that the N- and C-terminal fragments of Sca2 may interact with actin in a similar manner. To test this idea, we used ITC to assess the interaction of the full-length passenger domain of Sca2 (construct Sca_34-1515_) with monomeric actin. Latrunculin B (LatB), a marine toxin that binds in the catalytic site near the nucleotide in actin, was used in these experiments to prevent actin polymerization. The titration of LatB-actin into Sca_34-1515_ produced an exothermic reaction (Fig. 2A), whose binding isotherm fitted best to a two-site binding model (i.e. two actin subunits per Sca_34-1515_ molecule), and both sites had high affinity (K_D1_ = 0.19 μM and K_D2_ = 0.02 μM) (Fig. 2A, blue curve). Fitting to a single site model, shown for comparison (Fig. 2A, red curve), produced a much worse χ^2^ value (109.5 compared to 8.5) and unrealistic stoichiometry (N = 0.6). Note that the χ^2^ values reported here are reduced, i.e. divided by DoF (degrees of freedom), and while these values provide a measure of the goodness of the fit, they cannot be compared across different experiments, because the data are not weighted and the χ^2^ depends on the magnitude of the scale.

**Figure 2.**
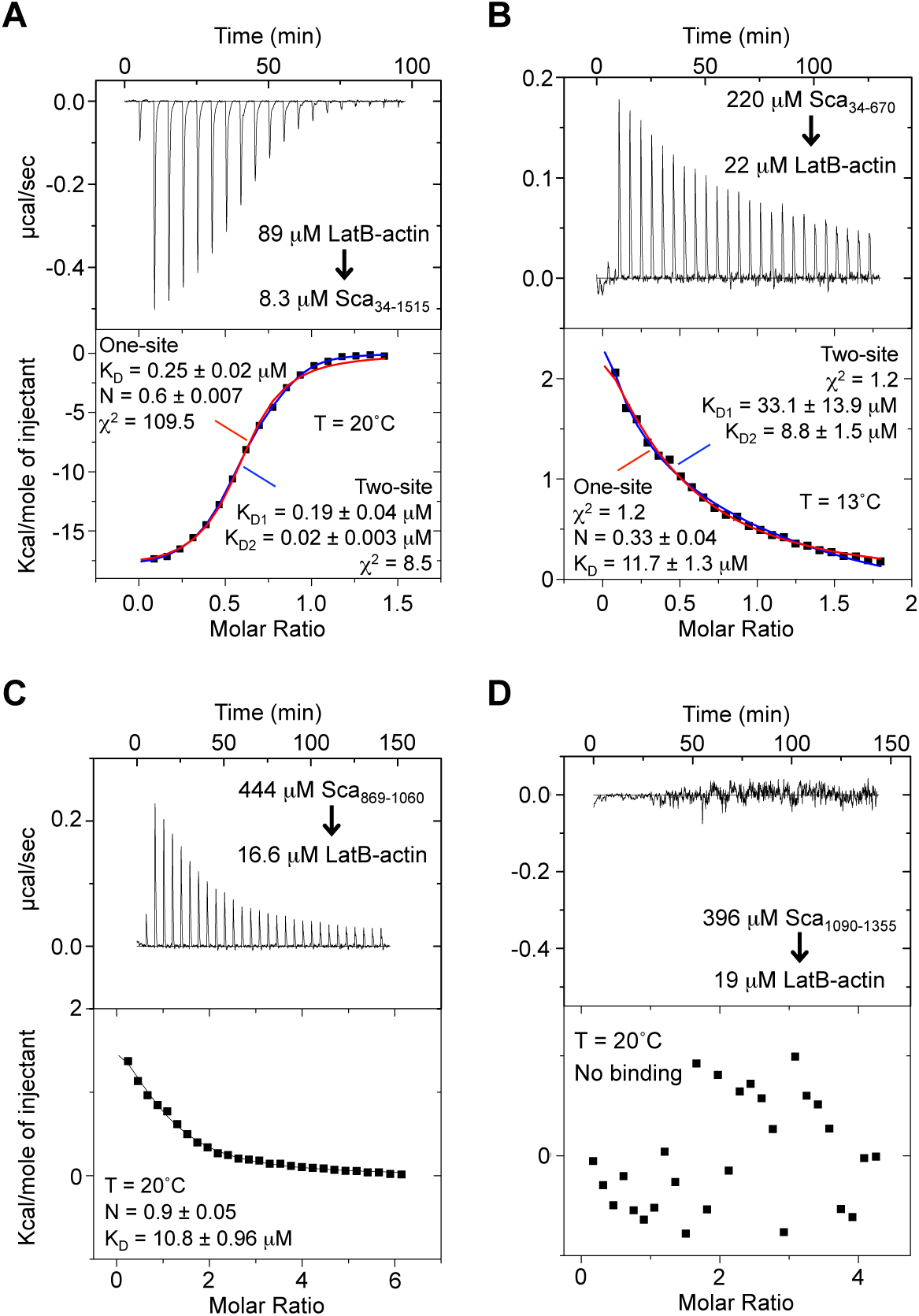
Interaction of Sca2 constructs with monomeric actin. A. ITC titration of 89 μM LatB-actin (in the syringe) into 8.3 μM Sca_34-1515_ (in the cell). The experiment was performed at 20°C. A two-site binding model (blue trace) produced a better fit of the data to a binding isotherm than a one-site binding model (red trace), as indicated by the χ^2^ values and realistic stoichiometry. The χ^2^ values reported here are reduced, i.e. divided by DoF (degrees of freedom). Note that while the χ^2^ value clearly defines the best fitting model for a particular titration, they are not generally comparable across different titrations, because the data are not weighted and the χ^2^ depends on the magnitude of the scale. The dissociation constant (K_D_) and binding stoichiometry (N) derived from the fits are listed. Errors correspond to the s.d. of the fits.
B. ITC titration of 220 μM Sca_34-670_ (in the syringe) into 22 μM LatB-actin (in the cell). The experiment was performed at 12.7°C. A two-site binding model (blue trace) produced an almost equally good fit of the data to a binding isotherm than a one-site binding model (red trace), as indicated by the stoichiometry of the interaction. A two-site binding model is also supported by the fact that this fragment of Sca2 has nucleation activity (12). The dissociation constant (K_D_) and binding stoichiometry (N) derived from the fits are listed. Errors correspond to the s.d. of the fits.
C. ITC titration of 444 μM Sca_869-1060_ (in the syringe) into 16.6 μM LatB-actin (in the cell). The experiment was performed at 20°C. The dissociation constant (K_D_) and binding stoichiometry (N) derived from fitting of the binding isotherm are listed. Errors correspond to the s.d. of the fits.
D. ITC titration of 396 μM Sca_1090-1355_ (in the syringe) into 19 μM LatB-actin (in the cell). The experiment was performed at 20°C. No binding is reported, as the experiment could not be fit to a binding isotherm.

We then asked whether the N- and C-terminal fragments of Sca2 independently contributed to this interaction. Curiously, the titration of the N-terminal half of Sca2 (Sca_34-670_) into LatB-actin produced an endothermic reaction. The titration was first fitted to a one-site binding model, but the stoichiometry of the interaction was unrealistically low (N = 0.33), suggesting that Sca_34-670_ interacts with two actin subunits (Fig. 2B).

Fitting to a two-site binding model yielded two binding sites of lower affinity than those of the full-length passenger domain (K_D1_ = 33.1 μM and K_D2_ = 8.8 μM), but the χ^2^ value (1.2) did not improve. A two-site binding model is preferred in this case because it addresses the low stoichiometry obtained with the one-site binding model, and explains why this fragment of Sca2 displays nucleation activity albeit lower than that of the full-length passenger domain (12).

We could not test binding of the entire C-terminal region (Sca_868-1515_) to LatB-actin due to the very low yields of this construct when expressed in isolation. Instead, we tested binding of two fragments that make up this construct (Sca_869-1060_ and Sca_1090-1355_). Sca_869-1060_, which contains a predicted repeat of three WH2 domains (see below), bound only one LatB-actin monomer with a K_D_ of 10.8 μM (Fig. 2C), and the reaction was endothermic in character, like for Sca_34-670_. In contrast, Sca_1090-1355_ did not appear to bind LatB-actin (Fig. 2D). Therefore, these results suggest that the two halves of Sca2 contribute to actin binding, analogous to the individual subunits of the FH2 dimer (25, 26), but only when combined together can they achieve high affinity binding of two actin monomers. The fact that the interactions change from endothermic for the individual fragments to exothermic for the full-length passenger domain and the affinities increase many folds suggests that additional interactions, presumably including actin-actin interactions, are involved in the formation of this ternary complex.

### Sca2 does not contain WH2 domains

As established above, Sca2 binds only two actin monomers, whereas the region containing the putative repeat of three WH2 domains (Sca_869-1060_) only binds one actin monomer and with much weaker affinity than the full-length passenger domain. All the WH2 domains analyzed by ITC so far bind actin with higher affinity than Sca_869-1060_, and their binding is consistently exothermic in character (28-30). These observations raise questions about the authenticity of the WH2 domains of Sca2. This is important, because it is still assumed that Sca2 contains WH2 domains (22), and it was the identification of putative WH2 domains based on sequence analysis that supported the initial hypothesis that Sca2 was a bacterial actin assembly factor, albeit curiously two different laboratories had proposed different definitions of the WH2 domains (11, 13).

We use here the latest definition of the WH2 domains of Sca2 (11) (Fig. 3A), which produces a better alignment with other known WH2 domains (Fig. 3B). Yet, the initial disagreement is understandable when one compares the sequence of the WH2 domains of Sca2 with other canonical WH2 domains. The WH2 domain is ~17 amino acids in length, and consists of an N-terminal α-helix that binds in the hydrophobic or target-binding cleft of actin, a loop and a C-terminal LKKV motif (22). All three of the WH2 domains of Sca2 lack at least one critical element of the consensus sequence. Thus, WH2a (residues 772-888) has a proline residue within the LKKV motif (22). WH2b (residues 943-960) lacks a basic residue at the N-terminus and key hydrophobic residues in the N-terminal α-helix of the domain. WH2c (residues 1004-1020) lacks the two basic residues of the LKKV motif. Importantly, mutations in the LKKV motif of classical WH2 domains disrupt binding to monomeric actin (31,32).

**Figure 3.**
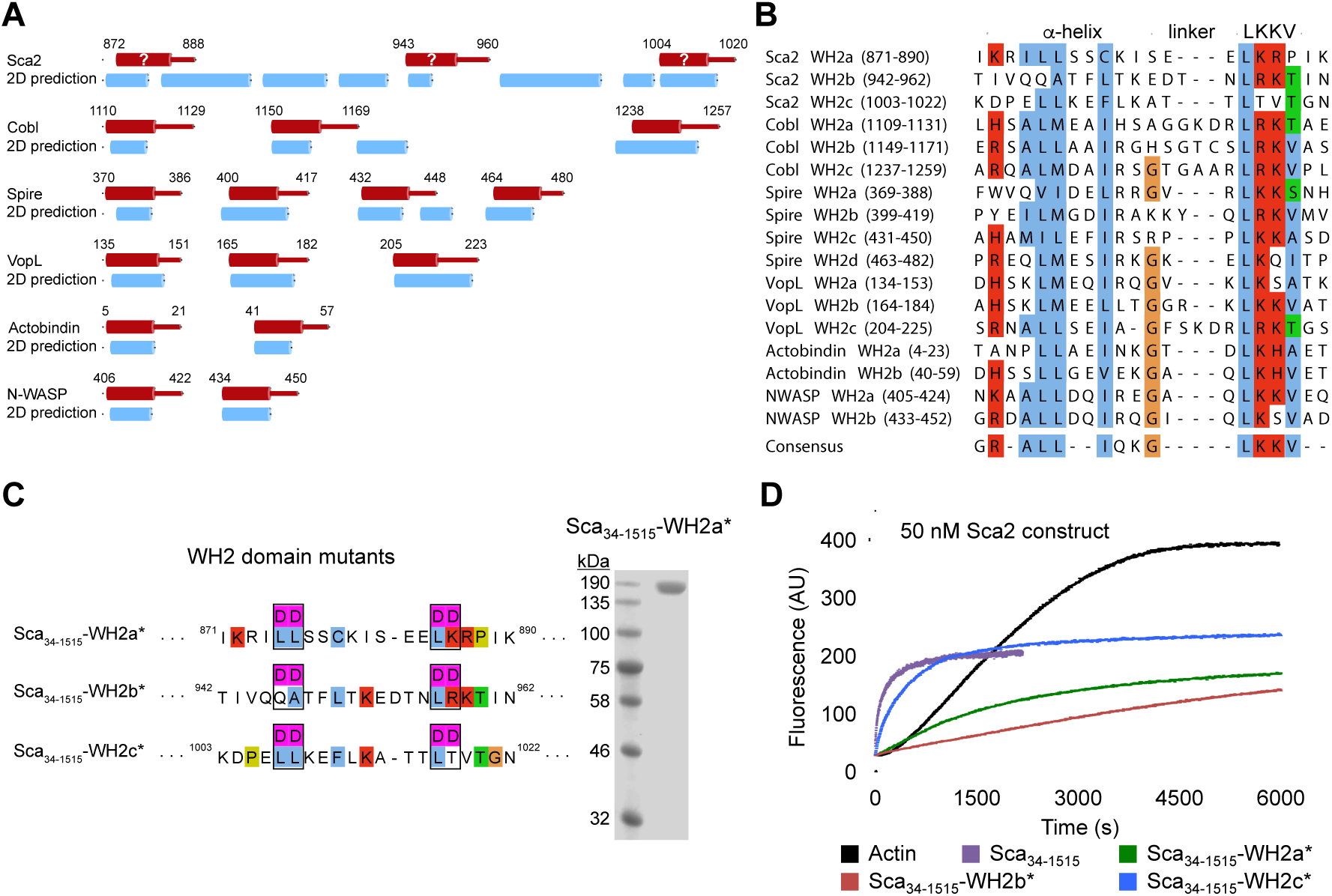
The region 868-1023 of Sca2 is critical for nucleation but lacks WH2 domains. A. Cartoon representation of the WH2 domain-containing regions of various cytoskeletal proteins containing tandem repeats of WH2 domains. WH2 domains are colored red, with a cylinder representing the α-helix of the domain. Secondary structure predictions obtained with the program Jpred (41) are also shown for each protein (2D prediction), with blue cylinders representing predicted α-helices.
B. Alignment of the sequences of the WH2 domains represented in part A, showing the regions corresponding to the N-terminal α-helix and LKKV motif, as well as the consensus sequence at the bottom.
C. Representation of the three mutants of Sca_34-1515_ (full-length passenger domain) targeting the putative WH2 domains. Each mutant carries four aspartic acid mutations of highly conserved residues of the canonical sequence of the domain. The SDS-PAGE shown on the right illustrates the purity of one of the mutants (Sca_34-1515_-WH2a^∗^).
D. Time course of polymerization of 2 μM Mg-ATP-actin (6% pyrene-labeled) alone and with the addition of 50 nM Sca_34-1515_ wild type and WH2 domain mutants depicted in part C (color-coded as indicated).

We then analyzed the sequences connecting the WH2 domains of Sca2. Here again, Sca2 is a clear outlier, with the putative WH2 domains spaced farther apart (45-55 amino acids) than in other proteins containing repeats of WH2 domains (~15 amino acids) (Fig. 3A). A flexible 15-amino acid linker seems to be optimal for proteins in which successive WH2 domains connect actin subunits along the long-pitch helix of the actin filament (33). The one exception is Cobl, where a 69-amino acid linker connects the second and third WH2 domains. However, Sca2 differs in yet another way from canonical WH2 domain-containing proteins; the sequences between WH2 domains are predicted to contain a series of α-helices, whereas in other proteins these sequences are typically unstructured (see the secondary structure predictions in Fig. 3A). Hydrophobic cluster analysis (HCA) further supports this conclusion, revealing a series of clusters of hydrophobic residues in the inter-WH2 domain linkers of Sca2 but not other proteins (Fig. S1). Such clusters typically denote the presence of secondary structural elements, which depending on the shape of the clusters can be defined as either α-helices or β-strands (34). The HCA plots of the WH2 domains themselves are also different between Sca2 and other WH2 domain-containing proteins (Fig. S1). The fact that the putative WH2 domains of Sca2 form part of a globular domain is finally supported by far-UV circular dichroism analysis of a Sca2 fragment comprising amino acids 868-1515, revealing a typical α-helical profile with minima at 208 and 222 nm (12).

Therefore, several lines of evidence suggest that Sca2 does not contain a typical WH2 domain repeat. However, we had previously found that a full-length passenger domain construct carrying four mutations in each of the WH2 domains (a total of 12 mutations targeting both the α-helix and LKKV motif of each WH2 domain) had no detectable nucleation or elongation activity (12), demonstrating the importance of this region in actin assembly. But, because this region binds only one actin monomer (Fig. 2C), we asked whether only one of the putative WH2 domains was actually functional. For this, we generated three Sca_34-1515_ mutants, individually targeting each of the WH2 domains. We used the same mutations as before (12), i.e. two conserved hydrophobic residues within
the N-terminal α-helix and the first two residues of the LKKV motif were simultaneously mutated to aspartic acid (Fig. 3C). Using the pyrene-actin polymerization assay, we found that the mutants targeting WH2 domains a and b, Sca_34-1515_-WH2a^∗^ and Sca_34-1515_-WH2b^∗^, inhibited the polymerization activity in a concentration-dependent manner, whereas mutant Sca_34-1515_-WH2c^∗^ had a negligible effect compared to the wild-type passenger domain (Fig. 3D and Fig. S2).

Therefore, at least two of the predicted WH2 domains of Sca2 are important for actin assembly within the full-length protein – is this because they bind actin directly the way WH2 domains do? To address this question, we generated four maltose-binding protein (MBP) fusion constructs, with the WH2 domains of Sca2 or a control WH2 domain from WAVE1 fused C-terminally to the MBP moiety, and separated by a non-cleavable 15-amino acid linker (Fig. 4A). The proteins were highly pure, and the yields were sufficiently high to allow for quantitative ITC experiments to assess their binding to LatB-actin (Fig. 4B). As anticipated, the first WH2 of WAVE1 (WAVE1 WH2a) bound LatB-actin with high affinity and 1:1 stoichiometry (K_D_ = 0.38 μM, N = 1.07) and the reaction was exothermic (Fig. 4C). This interaction is therefore very similar to that of synthetic WH2 peptides from other cytoskeletal proteins we have measured before in G-buffer (i.e. without the addition of LatB) or in F-buffer with the addition of LatB (28-30). In contrast, none of the putative WH2 domains of Sca2 bound LatB-actin (Fig. 4D-F), conclusively demonstrating that these are not functional WH2 domains.

**Figure 4.**
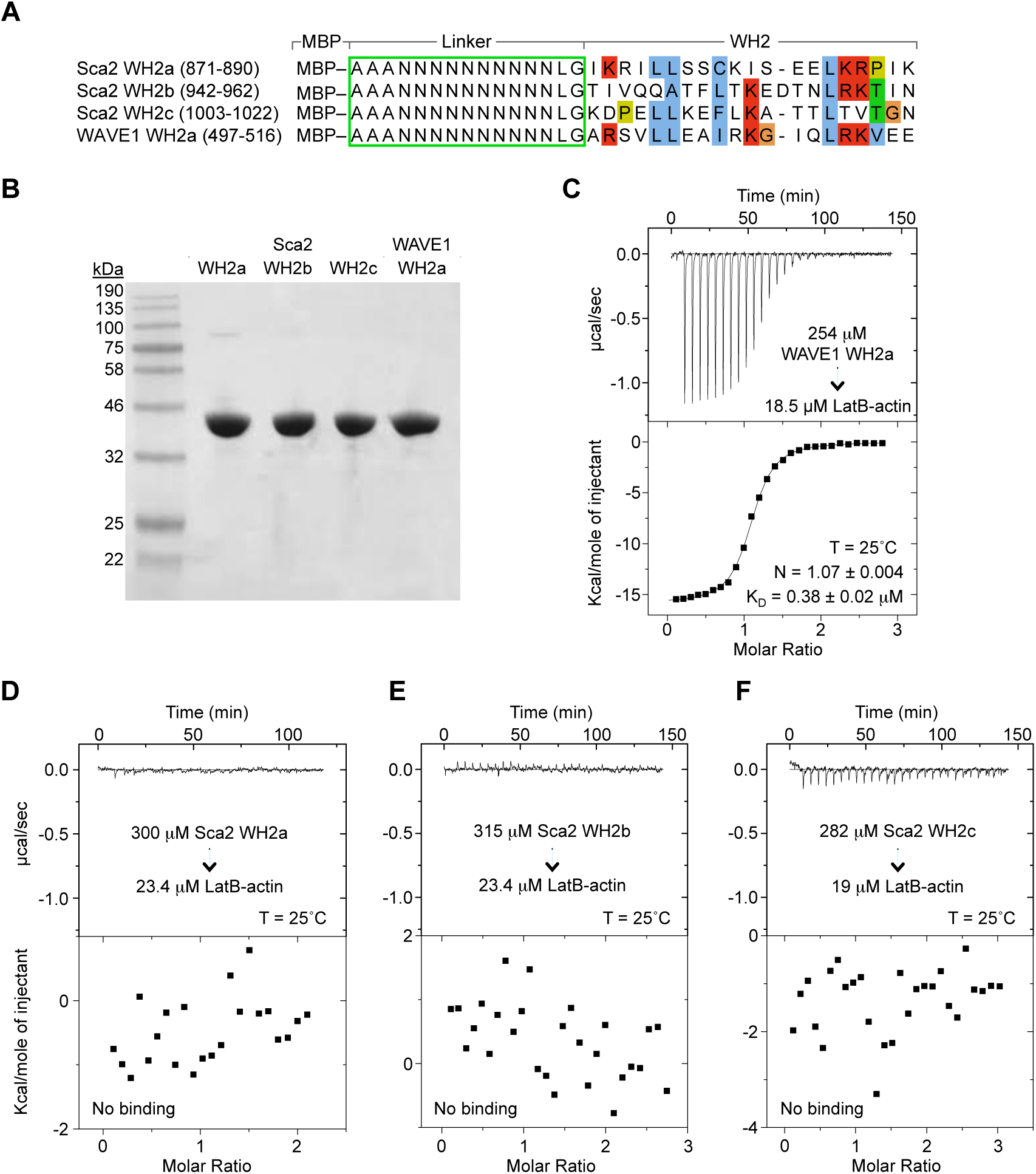
The putative WH2 domains of Sca2 do not bind actin. (A) Representation of four WH2 domain constructs expressed here as fusion proteins with maltose-binding protein (MBP). A 15-amino acid flexible linker separates the MBP moiety at the N-terminus from the WH2 domain of WAVE1 (control) or the three putative WH2 domains of Sca2.
(B) Purity of proteins illustrated in part A assessed by SDS-PAGE analysis.
(C) ITC titration of 254 μM WAVE1 WH2a into 18.5 μM LatB-actin. The experiment was performed at 25°C. The dissociation constant (K_D_) and binding stoichiometry (N) derived from fitting of the binding isotherm are listed. Errors correspond to the s.d. of the fits.
(D-F) ITC titrations of the three constructs of putative WH2 domains of Sca2 into LatB-actin. The concentrations of the proteins in the syringe and in the cell are indicated. The titrations could be fitted to a binding isotherm, as none of the putative WH2 domains of Sca2 appeared to bind LatB-actin.

## Discussion

*Rickettsia* Sca2 is a pathogenic bacterial actin assembly factor that mimics eukaryotic formins in that it generates long and unbranched actin filaments by combining both nucleation and processive barbed end elongation activities (11). However, we have shown before that Sca2 is structurally unrelated to formins and uses a different mechanism for actin assembly (12). Specifically, Sca2 is monomeric and lacks the dimeric FH2 domain found in all eukaryotic formins, which is crucial for both nucleation and elongation (35). The one element of the Sca2 sequence that still appeared to resemble eukaryotic nucleators was the postulated repeat of three WH2 domains (22), whose identification first drew attention to this protein as a potential actin assembly factor (11, 13). Of note, several eukaryotic formins contain WH2 domain-related sequences or form complexes with proteins that contain WH2 domains and synergize with formins for nucleation (36-40). Here, however, we have disproven the existence of WH2 domains in Sca2, suggesting that *Rickettsia* Sca2 has evolved completely independently of any eukaryotic actin assembly factor to acquire the various complex functions associated with eukaryotic formins: nucleation, elongation, and inhibition of barbed end capping (11, 12).

How does Sca2 accomplish these functions using a different fold? In thinking about this question, a comparison with eukaryotic formins provides some clues. Formins are dimeric, and specifically the two FH2 domains of the dimer interact head-to-tail to form a ring-like structure (24), which encircles two actin subunits at the barbed end of the actin filament while loosely contacting a third incoming subunit during processive elongation (23). Sca2 cannot be a dimer, since bacterial autotransporters are either monomeric or trimeric, depending on the fold of the translocator domain, which can be ~300 (monomeric) or ~75 (trimeric) amino acids long (18). The passenger domain of Sca2 is monomeric in solution (12) and, consistently, Sca2 has a 281-amino acid monomeric type of translocator domain. However, like formins, Sca2 must ‘hug’ the barbed end of the filament, while allowing for free access of additional actin subunits for processive elongation. This can only be accomplished if Sca2 also forms a ring-like structure around the barbed end, contacting primarily two actin subunits, and possibly a third incoming subunit (Fig. 5). The data presented here is consistent with this model. The N- and C-terminal regions of Sca2 (Sca_34-670_ and Sca_868-1515_) seem to behave to a large extent like the FH2 domains of the formin dimer: a) they bind actin individually but with weak affinity (Sca_34-670_ interacts with two actin subunits whreas Sca_868-1515_ appears to contact only one subunit), b) together (i.e. the full-length passenger domain) they bind two actin subunits with high affinity and account for the nucleation and barbed end elongation activities of Sca2, which the individual fragments mostly lack (12), c) deletions at either end of these two fragments disrupt their ability to interact with each other, suggesting that they bind head-to-tail to form a ring-like structure analogous to the formin FH2 domain. One of these fragments, Sca_869-1060_, comprises what was previously thought to be a repeat of three WH2 domains. However, Sca_869-1060_ binds a single actin subunit with weak affinity and following an endothermic reaction, clearly distinct from the higher affinity and exothermic reactions of canonical WH2 domains (28-30). Moreover, we have shown here that none of the putative WH2 domains binds actin in isolation, suggesting that the adverse effect of mutations of two of the domains is likely due to disruption of a contiguous actin-binding surface within this region of Sca2, which we now call the ‘middle domain’ because of its location within the full-length passenger domain (Fig. 5). Analogous to the proline-rich FH1 domain of formins, Sca2 also contains two proline-rich segments (672-699 and 1077-1087) that mediate the binding of profilin-actin for processive barbed end elongation (11, 12). Both segments form part of loop regions (658-714 and 1066-1091), which are predicted to be unstructured and could therefore be loosely attached to the other domains of the Sca2 molecule that form the ring-like structure around the barbed end. None of the Sca2 fragments studied here could be crystallized (except Sca_34-400_ which we studied previously (12)), either alone or in complex with actin, but structural studies testing some of the predictions of this model should soon become possible using cryo-electron microscopy.

**Figure 5.**
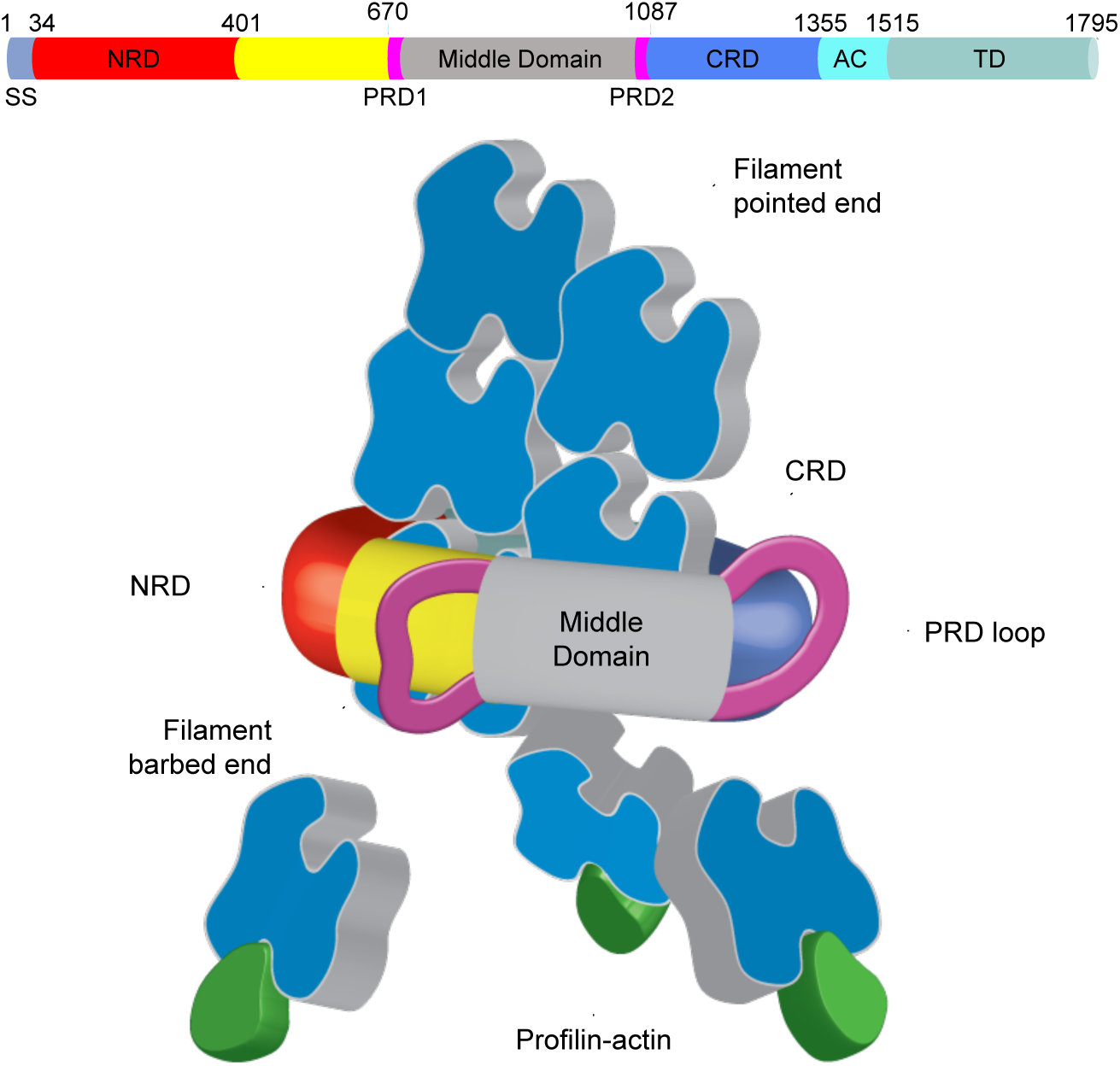
Proposed mechanism of nucleation and elongation by *Rickettsia* Sca2. By analogy with eukaryotic formins, Sca2 is proposed to form a ring-like structure around the barbed end of the actin filament, contacting primarily two actin subunits, and possibly a third incoming subunit. The data presented here supports this model, and suggests that the N- and C-terminal regions of Sca2 (Sca_34-670_ and Sca_868-1515_) behave in a way analogous to the FH2 domain of formins. Thus, we have shown here that these two fragments bind actin individually but with weak affinity, whereas the full-length passenger domain binds two actin subunits with high affinity and has both nucleation and elongation activities, whereas the individual fragments are mostly inactive (11, 12). Furthermore, deletions at either end of these two fragments disrupt their ability to interact with each other, suggesting that they interact head-to-tail to form a ring-like structure analogous to the formin FH2 domain. We have also shown that Sca2 does not contain WH2 domains. The region previously though to contain the repeat of three WH2 domains (Sca_869-1060_) binds actin as a whole, but only one actin subunit and in manner clearly distinct from that of canonical WH2 domains. Accordingly, this region is represented here as forming part of the ring-like structure of Sca2, and is renamed as the ‘middle domain’ because of its location in the sequence. The two PRDs, implicated in the recruitment of profilin-actin for barbed end elongation (12), may project out, since they form part of loops that are predicted to be unstructured.

## Author Contributions

S.S.A. and R.D. designed the experiments. S.S.A. and I.-G.L. prepared the proteins, including cloning, expression and purification, and S.S.A. performed all of the experiments for this work. S.S.A. and R.D. wrote the manuscript. All authors analyzed the data and reviewed the figures and manuscript.

## Acknowledgements

We would like to thank Grzegorz Rebowski for the preparation of actin for this study and help with Fig. 5, and Malgorzata Boczkowska for help with the actin polymerization assay, ITC experiments and data analysis. This work was supported by NIH grant R01 GM073791.

